# eDNA metabarcoding shows highly diverse but distinct shallow, mid-water, and deep-water eukaryotic communities within a marine biodiversity hotspot

**DOI:** 10.1101/2024.07.14.603441

**Authors:** Patricia Cerrillo-Espinosa, Luis E. Calderón-Aguilera, Pedro Medina-Rosas, Jaime Gómez-Gutiérrez, Héctor Reyes-Bonilla, Amílcar Cupul-Magaña, Ollin T. González-Cuellar, Adrian Munguia-Vega

**Affiliations:** Centro Universitario de la Costa, Universidad de Guadalajara, Puerto Vallarta, Jalisco, Mexico; Applied Genomics Lab, La Paz, Baja California Sur, Mexico; Departamento de Ecología Marina, Centro de Investigación Científica y de Educación Superior de Ensenada, Ensenada, Baja California, Mexico; Departamento de Plancton y Ecología Marina, Centro Interdisciplinario de Ciencias Marinas, Instituto Politécnico Nacional, La Paz, Baja California Sur, Mexico; Laboratorio de Sistemas Arrecifales, Universidad Autónoma de Baja California Sur, La Paz, Baja California Sur, Mexico; Laboratorio de Ecología Marina, Centro Universitario de la Costa, Universidad de Guadalajara, Puerto Vallarta, Jalisco, Mexico; Sociedad de Historia Natural Niparajá A.C., La Paz, Baja California Sur, Mexico; Conservation Genetics Laboratory, School of Natural Resources and the Environment, The University of Arizona, Tucson, Arizona, USA

## Abstract

As the impact of human activities continues to move beyond shallow coastal waters into deeper ocean layers, it is fundamental to describe how diverse and distinct the eukaryotic assemblages from the deep layers are compared to shallow ecosystems. Environmental DNA (eDNA) metabarcoding has emerged as a molecular tool that can overcome many logistical barriers in exploring remote deep ocean areas. We analyzed thirty-two paired seawater samples collected via SCUBA and Niskin samplers from shallow (< 30 m) and deep ecosystems (40-500 m) within a recognized hotspot of marine biodiversity (Gulf of California, Mexico). We sequenced an eDNA metabarcoding library targeting the COI gene of eukaryotes. We demonstrated that the diversity of operational taxonomic units (OTUs) did not peak at shallow coastal regions and that the deep samples had similar levels of biodiversity to shallow sites but detected a significant vertical zonation between shallow and deeper habitats. Our results suggest that the deep refugia hypothesis applies to about a third of the 5495 OTUs identified that were shared between shallow and deep layer samples. At the same time, most taxa were exclusive from either shallow or deep zones. The observation that deep communities were as rich but quite distinct as shallow communities supports extending spatial management and conservation tools to deeper habitats to include a significant fraction of phylogenetic and functional diversity exclusive of mid and deep-water ecosystems.

## Introduction

As the footprint of human activities continues to expand in the tridimensional space of the oceans (Halpern et al., 2019), the attention to natural resources present in the ocean depths also has increased, including the mid-water layer between 30-150 m depth and the deep-water ocean beyond 200 m depth. Interest in deeper regions of the oceans originate from abundant fish resources (Irigoien et al., 2014, Pham et al., 2014), an important role in nutrient regeneration and biochemical processes to sustain the ocean’s productivity throughout the biological pump (Martin et al., 2020), gas and oil exploration (Cordes et al., 2016), deep-seabed mineral mining (Flora, 2023), bioprospecting and biomimetics commercial applications (Blasiak et al., 2022) and the potential impacts of ocean-based climate interventions (Levin et al., 2023). The effective management and conservation of marine ecosystems and their ecological services require knowing which species are present in them, and their distribution in time and space. Our understanding of marine biodiversity and the impact of human activities has historically focused on shallow (< 30 m) coastal waters (Webb et al., 2010; Bongaerts et al., 2019), while mid-water depths and deep-water ecosystems have been considerably less studied (Eyal et al., 2021; Jacquemont et al., 2024). Multiple logistical constrains explain the lag in describing biodiversity beyond shallow waters, including the limited sampling accessibility, expertise and prohibitive costs of some of the most common exploration tools like deep-sea submersibles, remotely operated vehicles (ROVs) or sensors (Bell et al., 2023). One promising technology that could boost exploration of biodiversity in the ocean depths is environmental DNA (eDNA) metabarcoding where a sample of seawater or sediment taken remotely (e.g. with a water or sediment sampler) is processed to capture, amplify and massively sequence a conserved genomic region or metabarcode and used to detect the presence and biodiversity of taxa within a sample (Sinniger et al., 2016; Thomsen et al., 2016).

A key scientific question is how rich and distinct the biological communities from the mid-water depths, and deep-water are compared to shallow ecosystems. A global meta-analysis suggested species diversity in the ocean decreases with depth, and that the 0-100 m depth range contains up to four times the diversity recorded between 100-200 m (Costello & Chaudhary, 2017). The deep refugia hypothesis suggests that mid-water depth reefs could act as a refugee from disturbances for shallow reef communities, and implicitly assumes that many species show wide depth ranges and considerable vertical ecological connectivity between them (Riegl & Piller, 2003; Bongaerts et al., 2010) sometimes avoiding regional extinction of particular species (del Monte-Luna et al., 2023). Multiple observational studies in fish have shown that mid-water depth communities are diverse but taxonomically and functionally distinct from their shallow counterparts (Rocha et al., 2018, Medeiros et al., 2021; Loiseau et al., 2022), while evidence supporting vertical connectivity within fish species occurring at different depths due to diel vertical migration is mixed (Tenggardjaja et al., 2015; Loya et al., 2016). Studies on benthic communities have shown strong vertical zonation in function of seafloor depth, low connectivity, and a clear distinction between shallow and mid-water depth benthic communities (Bongaerts et al., 2017, Stefanoudis et al., 2019).

Our study focused in the Gulf of California (*Fig. 1*), a globally recognized hotspot of marine biodiversity on the Northwest of Mexico that is ∼1,500 km long, ∼100 km width covering 12 degrees of north latitude and characterized by seasonally reversing ocean gyres that sit on deep basins reaching up to 4 km deep (Munguia-Vega et al., 2018). The Gulf of California is a highly productive tropical-subtropical system that supports more than half total Mexico’s marine fisheries and an economically profitable growing ecotourism industry, but nonetheless shows signs of significant ecosystem decline due to overfishing and climatic change (Gilly et al., 2022). A recent eDNA metabarcoding study showed that biodiversity levels from shallow coastal areas in the Gulf of California are much higher than previously assumed based on historical records and visual surveys (Mac Loughlin et al., 2024), but few studies exist on the biota from mid-water depths and deep-water ecosystems. These studies have focused on the central Gulf of California (Figure 1a) based on fish ROV surveys from the mid-water depth zone (Hollarsmith et al., 2020, Velasco-Lozano, 2020) and ROV and submersible fish and invertebrate surveys of the deep-waters (Aburto-Oropeza et al., 2011; Portail et al., 2016, Gallo et al., 2020).

**Figure 1 a, b:**
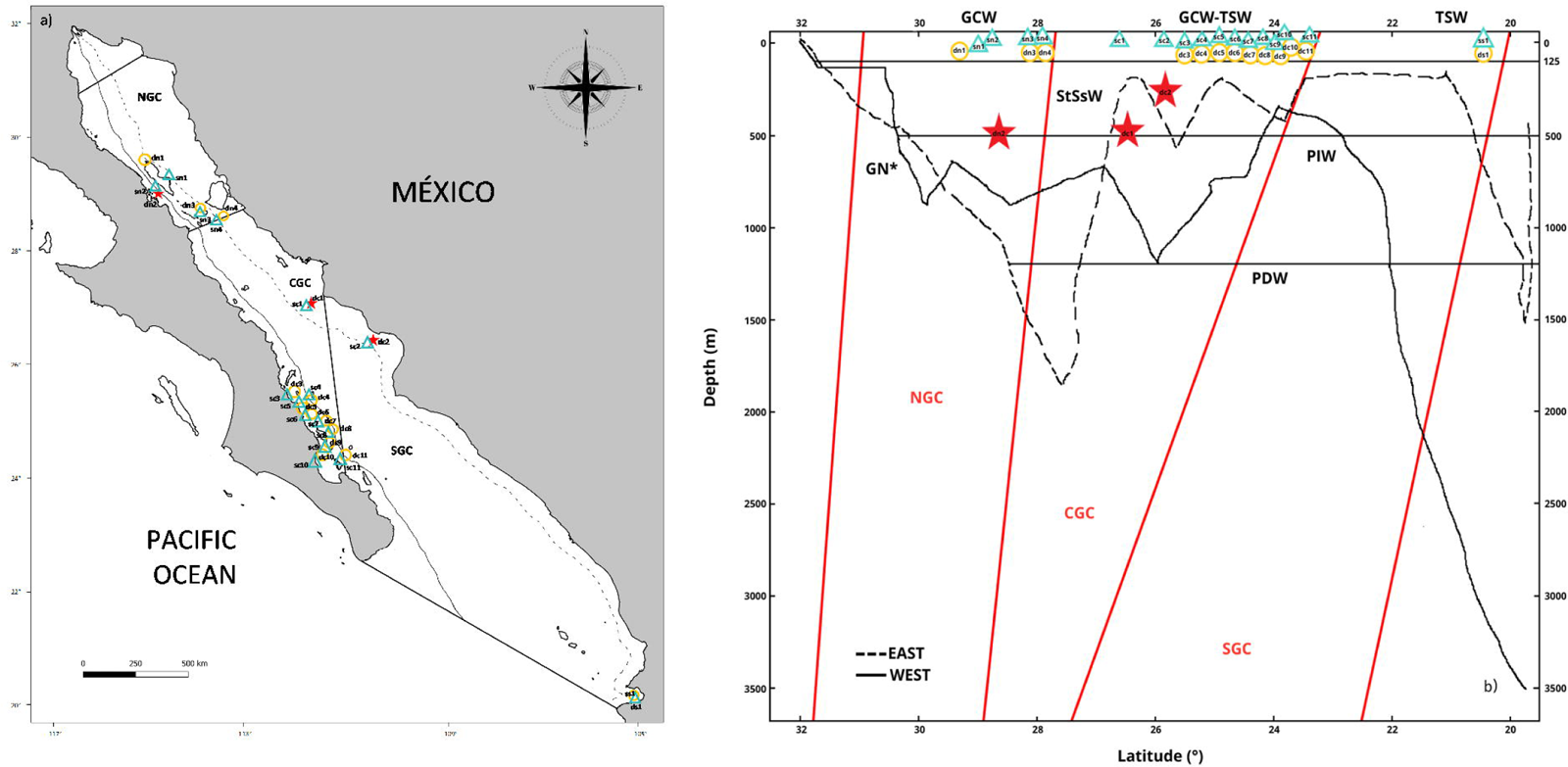
Sampling Sites of the Study Area. Sampling sites distributed across the three biogeographic regions including the Northern (NGC), Central (CGC) and Southern (SGC) Gulf of California (a). Sampling sites according to vertical gradient, sea water masses (horizontal lines), and biogeographic regions (red lines) (b). Shallow sites are in green color, mid-water depth sites are in yellow, and deep-water sites are indicated as red stars. See Table 1 for the classification of water mass for each sample.

**Table 1:**
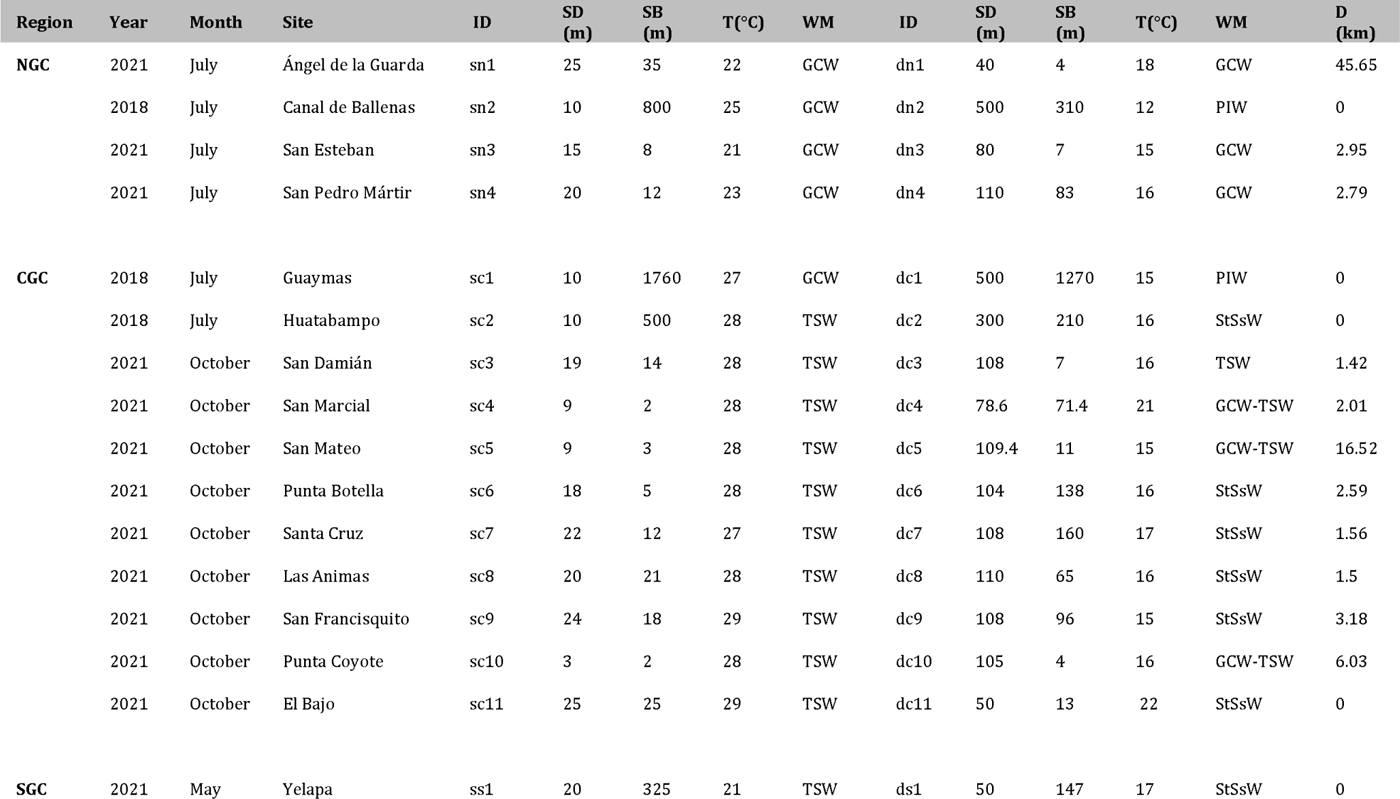
Sampling data per region, including sampling date, site and sample ID. SD= Sample depth, SB= Sample site distance to bottom, D= Distance between shallow and deep sites, T= Temperature. WM= Water masses follow Lavín & Marinone (2003): GCW= Gulf of California Water; TSW= Tropical Surface Water; StSsW= Subtropical Subsurface Water; PIW= Pacific Intermediate Water.

| Region | Year | Month | Site | ID | SD<br>(m) | SB<br>(m) | T(°C) | WM | ID | SD<br>(m) | SB<br>(m) | T(°C) | WM | D<br>(km) |
| --- | --- | --- | --- | --- | --- | --- | --- | --- | --- | --- | --- | --- | --- | --- |
| NGC | 2021 | July | Ángel de la Guarda | sn1 | 25 | 35 | 22 | GCW | dn1 | 40 | 4 | 18 | GCW | 45.65 |
|  | 2018 | July | Canal de Ballenas | sn2 | 10 | 800 | 25 | GCW | dn2 | 500 | 310 | 12 | PIW | 0 |
|  | 2021 | July | San Esteban | sn3 | 15 | 8 | 21 | GCW | dn3 | 80 | 7 | 15 | GCW | 2.95 |
|  | 2021 | July | San Pedro Mártir | sn4 | 20 | 12 | 23 | GCW | dn4 | 110 | 83 | 16 | GCW | 2.79 |
| CGC | 2018 | July | Guaymas | sc1 | 10 | 1760 | 27 | GCW | dc1 | 500 | 1270 | 15 | PIW | 0 |
|  | 2018 | July | Huatabampo | sc2 | 10 | 500 | 28 | TSW | dc2 | 300 | 210 | 16 | StSsW | 0 |
|  | 2021 | October | San Damián | sc3 | 19 | 14 | 28 | TSW | dc3 | 108 | 7 | 16 | TSW | 1.42 |
|  | 2021 | October | San Marcial | sc4 | 9 | 2 | 28 | TSW | dc4 | 78.6 | 71.4 | 21 | GCW-TSW | 2.01 |
|  | 2021 | October | San Mateo | sc5 | 9 | 3 | 28 | TSW | dc5 | 109.4 | 11 | 15 | GCW-TSW | 16.52 |
|  | 2021 | October | Punta Botella | sc6 | 18 | 5 | 28 | TSW | dc6 | 104 | 138 | 16 | StSsW | 2.59 |
|  | 2021 | October | Santa Cruz | sc7 | 22 | 12 | 27 | TSW | dc7 | 108 | 160 | 17 | StSsW | 1.56 |
|  | 2021 | October | Las Animas | sc8 | 20 | 21 | 28 | TSW | dc8 | 110 | 65 | 16 | StSsW | 1.5 |
|  | 2021 | October | San Francisquito | sc9 | 24 | 18 | 29 | TSW | dc9 | 108 | 96 | 15 | StSsW | 3.18 |
|  | 2021 | October | Punta Coyote | sc10 | 3 | 2 | 28 | TSW | dc10 | 105 | 4 | 16 | GCW-TSW | 6.03 |
|  | 2021 | October | El Bajo | sc11 | 25 | 25 | 29 | TSW | dc11 | 50 | 13 | 22 | StSsW | 0 |
| SGC | 2021 | May | Yelapa | ss1 | 20 | 325 | 21 | TSW | ds1 | 50 | 147 | 17 | StSsW | 0 |

We analyzed paired water samples collected via SCUBA and Niskin samplers from shallow (< 30 m) and deep ecosystems (40-500 m) at the three biogeographic regions of the Gulf of California through eDNA metabarcoding of a fragment of the COI gene targeting eukaryotes. Using eDNA of eukaryotes we tested the deep refugia hypothesis that states that deep reef ecosystems, particularly those in the mid-water depths (>30 meters depth), could serve as refuges for species from troubled shallow reefs, offering protection from disturbances affecting shallower areas like climate change and human activity. Our goals were: 1) to contrast the diversity of eukaryotes between shallow and mid-water depth and deep-water layers of the Gulf of California, and 2) to establish how distinct the biological communities are according to vertical depth. Evidence on the diversity and distinctiveness of each vertical layer in the pelagic ecosystem could lead to a re-assessment of management and conservation priorities that traditionally have been mostly focused only on shallow ecosystems.

## Materials & Methods

### Sampling design

The study area is the Gulf of California, a marginal semi-closed sea of the north-eastern Pacific Ocean that has been recognized with three clearly defined biogeographic regions (North, Central and South, *Fig. 1a*) (Morzaria-Luna et al. 2018). Due to its oceanic connection to the Eastern Tropical Pacific and its location between temperate and tropical biogeographic regions, it is influenced by at least six sea water masses, each characterized by specific ranges of salinity, temperature and depth (*Fig. 1b*). The Pacific Deep Water (PDW) is distributed below 1,200 m depth; the Pacific Intermediate Water (PIW) below 500 m depth; the Subtropical Subsurface Water (StSsW) below the 150 m, the Tropical Surface Water (TSW) found at the surface at the Southern Gulf of California, the Gulf of California Water (GCW) found at the surface in the Central and Northern Gulf of California and formed by evaporation of StSsW and TSW and the modified California Current Water (CCW) which is present only in the Southern Gulf of California (Lavin & Marinone 2003; Monreal-Jiménez et al., 2021). We collected sea water samples at 16 sites distributed along the three biogeographic regions of the Gulf of California. Within each site, the experimental design included a shallow sample (< 25m) and a paired deep sample from either the mid-water depth (40 to 150 m) or deep-water layer (150 m to 500 m) (*Fig. 1b*, *Table 1*). Paired samples were collected from live aboard diving vessels (D/V Quino el Guardian) or oceanographic cruises (CAPEGOLCA, R/V El Puma).

Each shallow sample (n =16) from 0-30 m was paired either with a mid-water depth sample (n = 13) from 30-150 m or a deep-water sample (n = 3) from 150-500 m deep to obtain the eukaryotic eDNA signature at different column water depths within the same area and considering a mean general distance between sampling sites of 2.7 km (range 0 to 45 km, Table 1). The vertical position of the seawater sample in the water column was also examined by measuring the distance between the sampling depth to the bottom, following GEBCO bathymetry (GEBCO Compilation Group, 2021) (*Table 1*).

Shallow seawater samples were collected SCUBA diving above a depth of 25 m with Nalgene TM (Thermo Scientific) lab-quality broadmouth 1 L bottles (6 L total collected at different times during the dive at each site). A Niskin bottle with a capacity of 6 L operated by hand or attached to an oceanographic rosette was used to sample seawater at mid-water depth and deep-water depths between 40 and 500 m. We filtered 2 L of seawater at shallow and deep sites through the same filter for a total of three field replicates at each site and depth. Seawater samples were filtered with an electric pressure pump and nitrocellulose Millipore filters with a pore size of 0.45 μm placed in a Millipore Sterifil filter unit. Each filter was deposited in a 15 ml falcon tube with silica during field work (Miya & Sado, 2019) and refrigerated at 8°C back in the lab until processing. All the collecting and filtering equipment was cleaned by submersion in 1% sodium hypochlorite solution between sampling events and rinsed thoroughly with running freshwater. On every sampling day, 2 L of running fresh water used for cleaning the sampling and filtering equipment was collected and filtered on the field as a field control to test for external contamination.

### DNA extraction

Total DNA from environmental samples and negative field controls were extracted from the nitrocellulose filters with the DNeasy Blood & Tissue kit (QIAGEN,) according to the manufacturer’s instructions and using a QIAvac 24 Plus vacuum manifold to minimize contamination and handling. A blank negative control was incorporated at each extraction event. Total DNA concentration was measured for each sample with the Qubit 3.0 fluorometer (Invitrogen) and the High Sensitivity (Invitrogen) assay. All eDNA extractions were performed in a dedicated eDNA room and inside a hood used solely for this purpose. The hood and all the equipment and materials were sterilized with 1% sodium hypochlorite solution and UV light for 20 min between extraction events. Filter tips were used in all pipetting to reduce the risk of cross-contamination among seawater samples.

### Library preparation and sequencing

Partial sequences (313 bp) of the cytochrome oxidase subunit I (COI) barcode were amplified per triplicate for each individual DNA extraction (i.e.., nine PCR1 for each shallow and deep layer per site, respectively) with primers mICOIintF-XT: 5 GGWACWRGWTGRACWITITAYCCYCC 3 (Wangensteen et al., 2018) and dgHCO2198: 5 TAIACYTCIGGRTGICCRAARAAYCA 3 (Geller et al., 2013). The primer sets contained a standard Illumina adapter and an anchoring site for PCR2 primers following a library design provided previously (Valdivia-Carrillo et al., 2021). The amplification protocol for a 12 μl volume reaction included: 5 μl of eDNA (≥ 2ng), nuclease-free water, PCR Buffer (1X), MgCl2 (2.5mM), dNTP’s (0.2mM), Primers (0.4 μM each), 0.02% of BSA and 2 U of Platinum Taq HiFi polymerase (Invitrogen). PCR conditions were 95°C for 5 min, followed by 35 cycles of denaturation at 95°C for 30s, annealing at 45°C for 30 s, an extension of 72°C for 30 s and a final extension of 72°C for 5 min. PCR negative controls were included in each PCR. A simulated mock community constructed from equimolar concentrations of DNA from 25 known species of fish and invertebrates from six different phyla from the Gulf of California was used as a positive control and is described in detail previously (Mac Loughlin et al., 2023). Amplification of final products was verified in 1.2% agarose gels stained with RedGel (Biotium).

Individual barcode combinations for each seawater sample were introduced during PCR2 that were conducted in triplicate for all field samples, the mock community, three pools of field controls, three pools of DNA extraction controls, and three pools of PCR controls. The protocol for a final volume of 12 μl was: 3 μl from PCR1 pool, nuclease-free water, PCR Buffer (1X), 2.5mM of MgCl 2, 0.2mM of dNTP’s, 0.4 μM each primer, 0.02% BSA and 1 U Platinum Taq HiFi polymerase (Invitrogen). The thermocycling protocol was as follows: 95°C for 5 min, 12 cycles of 95°C for 30 s, 45°C for 30 s, 72°C for 15 s, and a final extension of 72°C for 5 min. The estimated size of the amplicon (448 bp) was verified in 1.2% agarose gels. The PCR2 products were quantified pooled for each individual sample for purification with AmpureXP beads (1.8X) (Beckman Coulter). The final products were quantified with Qubit and standardized to equimolar concentrations. The high-throughput sequencing of the library was carried out on the Illumina MiSeq platform (250bp x 2) at the University of Arizona Genetics Core.

### Sequence analysis

The bioinformatic analysis was performed in a Linux Ubuntu system v.20.04.1 (Sobell, 2015) using the USEARCH v11 software (Edgar 2010). Raw demultiplexed sequence reads were merged by maximum (380 bp) and minimum (280 bp) lengths where short alignments (<16 bp) were discarded, along with forward and reverse primers. The reads quality filter was done under a maximum expected number of errors 1.0. The reads were dereplicated with a minimum size (2 reads) to get the unique sequences and subsequently clustered (97% similarity threshold) into Operational Taxonomic Units (OTUs) using the UPARSE algorithm (Edgar, 2013), including detection and exclusion of chimeras. The last step consisted of the generation of the OTU table. The final OTUs were compared with the BLAST algorithm to the NCBI platform (Benson et al., 2013).

### Taxonomic assignments

Taxonomic assignments for each OTU were evaluated using the BLAST with the algorithm matching highly similar sequences. We generate XML files of the first one hundred results obtained for each OTU. The XML files were read in the MEGAN 6 Community Edition software (Huson et al., 2016) with parameters: Min score of 50.0, Min Percent Identity of 70.0, and Min Support Percent of 0.01. MEGAN used the Tree of Life from NCBI, the Last Common Ancestor algorithm (LCA, 100% to cover and the naive approach). Each OTU was statistically assigned to the LCA in the taxonomic tree, where the less consistency of taxonomic assignment, the higher up in the tree the assignment is placed for the OTU until the LCA of all likely assignments is reached. The taxonomic assignments were manually checked to discard cross-sample contamination and remove bacteria and terrestrial taxa. OTUs with no hits and no assignments in NCBI were compared against the BOLD Systems platform (Ratnasingham & Hebert, 2007) with the following similarity threshold: 100-97% (Species), 97-94% (Genus), 94-91% (Family), 91-88% (Order), 88-85% (Class) and <85% - >70% (Phylum) following Valdivia-Carrillo et al. (2021). A total of 1586 reads were assigned to 188 OTUs in the nine negative control samples from the field, extraction, and PCR steps, and these OTUs were eliminated from the entire dataset before statistical analyses.

### Statistical analysis

Histograms to explore the distribution of OTUs per sample were created with the ggplot2 package and Venn Diagrams with the Venn Diagram package in RStudio v2022.02.0 (R Studio Team, 2020). Species richness was estimated with Chao 1 non-parametric estimator based on the abundance of rare species using the Primer v7 software (Clarke & Gorley, 2015). We only considered the seawater samples from the North and Central regions of the Gulf of California to analyze the richness (alpha diversity) and test for significant differences between biogeographic regions. We found differences in OTUs richness sample size and therefore used the Mann-Whitney test. We used the Wilcoxon test to search for significant differences in OTUs richness between shallow and deep-sea water samples. Graphics were generated in Primer v7, and the statistical analyses were performed in XLSTAT software (Lumivero, 2023). We used the Jaccard presence/absence similarity resemblance matrix and a 2D non-metric multidimensional scaling (nMDS) ordination analysis in Primer v7 to test for eukaryote community structure based on biographic region, column water depth, and sea water mass. A one-way ANOSIM test was run in all cases with 9999 permutations, including one which considered the three variables (Biogeographic region, column water depth, sea water mass) to test the statistical significance. A permutational analysis of variance (PERMANOVA) was conducted in the PERMANOVA+ package from Primer v7 to analyze the community structure (beta diversity). Datasets were transformed into a presence/absence matrix, and the Jaccard test was applied to each comparison. We designed a 3-factor Global PERMANOVA (Biogeographic region, column water depth, and sea water mass) from the resemblance matrices. A pairwise analysis was additionally done with three, three, and four levels, respectively, based on expectations of mean squares: The biogeographic region (North, Central, South), column water depth (Shallow-Mid-water depth, Shallow-Deep-water, Mid-water depth-Deep-water), and sea water masses (GCW-TSW, GCW-PIW, GCW-StSsW, TSW-PIW, TSW-StSsW, PIW-StSsW). All the analyses were performed with 9999 permutations. To assess which of the taxa were responsible for the differences observed among biogeographic regions, column water depths, and sea water masses, we performed a two-way similarity percentage (SIMPER) survey (70% similarity threshold) in Primer v7, which was based on the presence/absence resemblance matrix and the Bray Curtis measure (relative frequencies). Finally, a Spearman correlation rank analysis was performed in XLSTAT to assess the relationship between richness and column water depth.

## Results

The sequenced library resulted in 4,665,588 total paired reads for the three column water depths, including controls (data deposited in GenBank Bioproject ID PRJNA1073001), with an average of 144,706 raw reads per sample (excluding controls) (*Table S1*). The USEARCH pipeline removed 2,270,370 reads during the merge step and 159,703 through quality control. The clusters <2 sizes were discarded (298,412 reads), along with 747,243 singletons and 31,775 chimeras. An OTU table was constructed from 2,110,667 reads, resulting in 228,953 unique reads grouped into 11,922 OTUs. The negative controls resulted in a total of 1,586 reads (*Table S2*), represented mostly by bacteria and the phyla Apicomplexa, Amoebozoa, Arthropoda, Mollusca, Cnidaria, Rhodophyta, Bacillariophyta and OTUs with no taxonomic assignment. We discarded a total of 6,427 OTUs, including those assigned to bacteria (2,468), terrestrial taxa (444), with no hits or taxonomic assignments (3,229) and all OTUs found in the negative controls (188, *Table S2*). The final analyses were conducted with 5,495 OTUs (*Table S3*). From these, 4,493 were taxonomically assigned with BLAST on the NCBI database, and 903 were assigned with the BOLD Systems platform. A total of 1,694 of these OTUs (30.8% from the total) were either assigned above the phylum taxonomic rank within Eukaryota, or these were taxa of the Stremenopiles, Alveolata, and Rhizaria lineages (SAR) for which higher level phylum taxonomy is still unresolved. The mock sample display a total of 4,065 reads grouped in 200 OTUs. Within the observed mock community, we successfully identified 20 species (80%) across various phylogenetic levels. Out of the expected 25 species, we detected 2 (9%) at the species level, 2 (9%) at the genus level, 5 (22%) at the family level, 4 (18%) at the order level, and 7 (30%) at the class level. We observed wide variation in the number of OTUs and reads assigned to each taxa within the mock community (*Table S4*).

We detected a total of 41 eukaryotic phyla among the samples analyzed. The five most diverse phyla in terms of OTUs were (in decreasing order): Arthropoda, Bacillariophyta, Mollusca, Rhodophyta and Cnidaria (*Fig. 2*). Most phyla showed a portion of OTUs that were unique to shallow or deep-sea water samples, and about one third of OTUs were found in both. Most phyla that were exclusively present in shallow samples (Evosea, Euglenozoa, Blastocladiomycota, Phoronida, Perkinsozoa, and Prasinodermophyta), or exclusive of deep-sea water samples (Heterolobosea, Chytridiomycota, and Mucoromycota) were represented by a single OTUs each one. Most of the taxonomically unassigned OTUs were exclusively found in the deep-sea water samples (705); followed by those OTUs shared between shallow and deep-sea water samples (531) and 458 OTUs were only found in the shallow depths.

**Figure 2:**
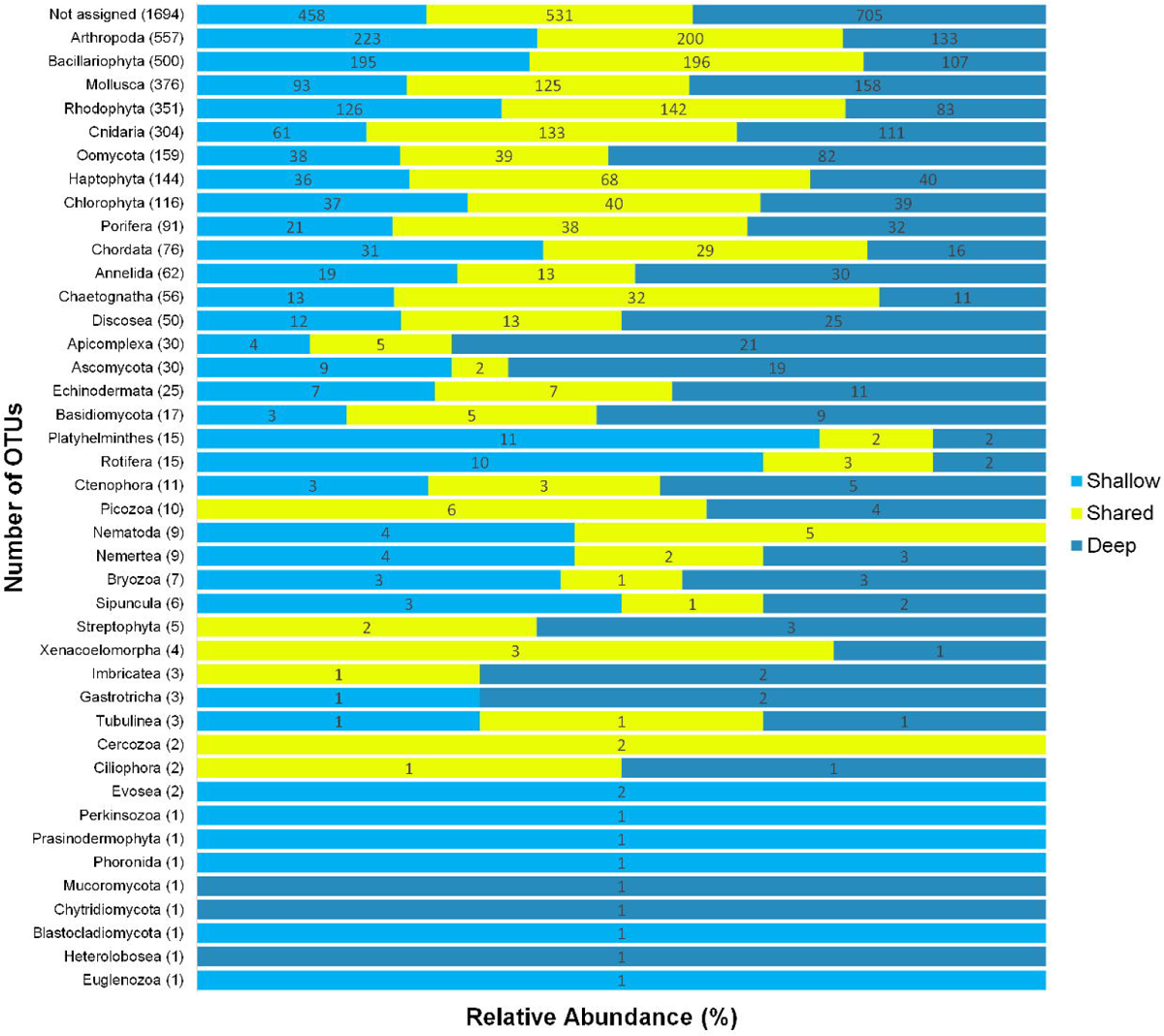
OTUs at a Phylum Level. OTUs counts at the phylum level from sea water samples collected at shallow and deep layers in the Gulf of California.

The Chao1 species richness estimator did not reach the asymptote for the total number of samples, particularly for the deep water samples, suggesting that the sampling effort was still insufficient to reach saturation of the OTUs present in these communities (*Fig. S1*). The combined OTUs richness from shallow and deep-sea water samples within sites averaged 893.4 OTUs (range = 272–1719) and was highly heterogeneous among the 16 sampling sites and within the three layers compared of each site (*Fig. 3*). Few sites with deep-water samples showed more than double the OTUs richness than their shallow counterparts (e.g., n1, c2, c3, and c6) and some shallow sites had the opposite trend (e.g., n2, n4, c1, c5, c7, and c11). Within all sampling sites, the cumulative richness observed was larger than that observed for the individual shallow or deep-water samples, indicating that a variable fraction of the OTUs within each site was not shared between shallow and deep water samples. The mean alpha diversity for the shallow sea water samples was slightly higher than the deep water samples (Shallow = 537.4; Deep = 481.0, *Fig. 4a*); but differences were not statistically significant (Wilcoxon *p* = 0.077). OTUs mean richness of the sites in the Northern Gulf of California were comparatively higher than those collected in the Central Gulf of California (Northern = 688.2; Central = 439.5, *Fig. 4b*), but differences were also not statistically significant (Mann-Whitney, *p* = 0.153), (Supplementary *Tables S5, S6*).

**Figure 3:**
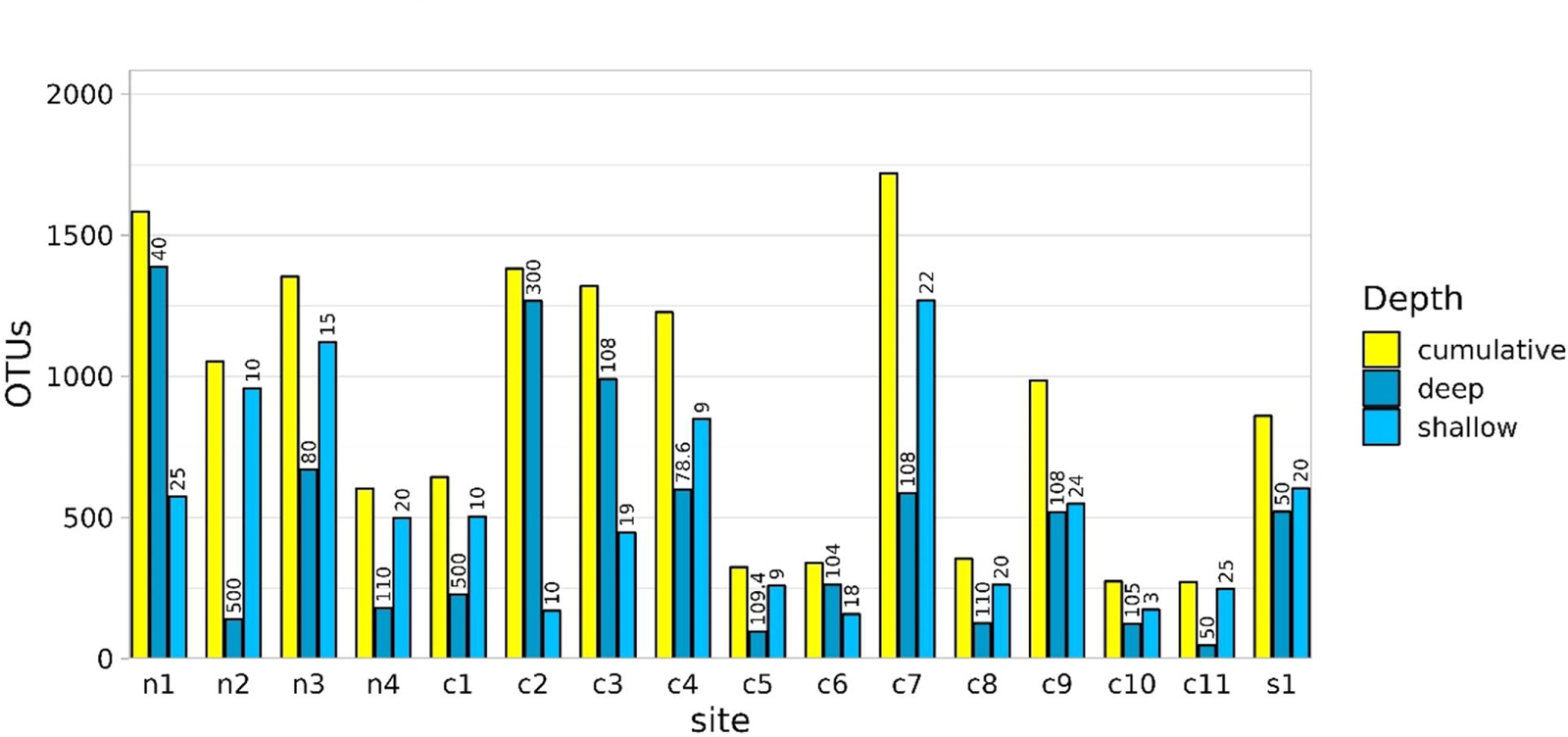
Species Richness. OTUs species richness per sampling depth ordered at latitudinal distribution (Northern samples on the left). The depth (m) from each sample is shown at the top of each bar.

**Figure 4 a,b:**
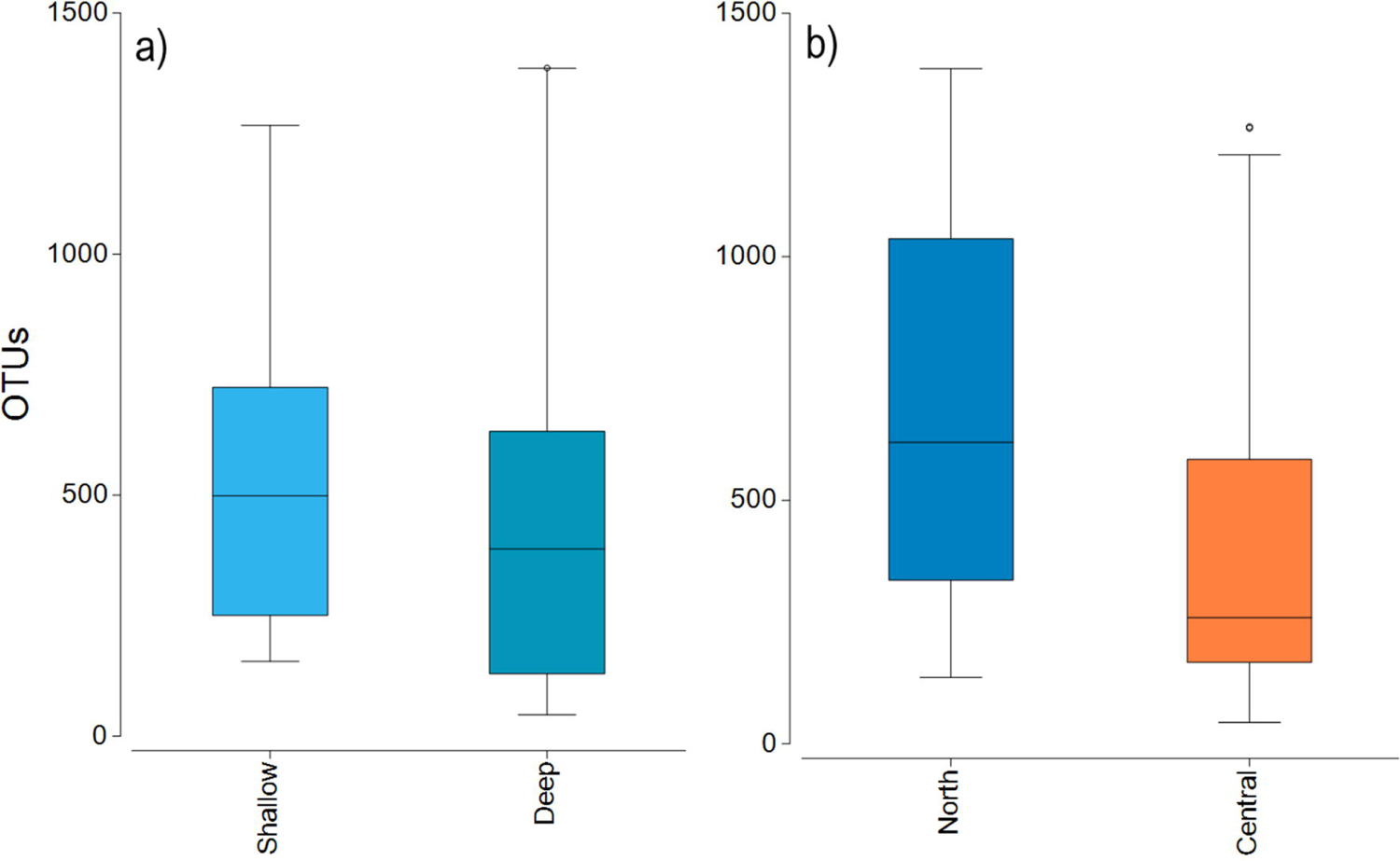
Alpha Diversity. Box plots showing the mean and 95% confidence intervals of OTUs species richness between shallow and deep-water samples (a) and between latitudinal regions (b).

About one third of the OTUs recorded in all sites from the Gulf of California were exclusive from shallow samples (30.8%), a third were exclusive from the deep-sea water samples (34.2%), and a third were shared between both (34.9%, *Fig. 5a*). The analyses of taxonomic assignments at Class level showed that most of Classes were shared between shallow and deep-sea water samples. However, 16 taxa from 12 different Phyla (Ciliophora, Pseudofungi, Amoebozoa, Platyherlminthes, Euglenozoa, Chordata, Bryozoa, Mollusca, Blastocladiomycota, Ascomycota, Basidiomycota and Chlorophyta) were exclusive from shallow sea-water samples and nine taxa from eight distinct phyla (Choanozoa, Mucoromycota, Platyhelminthes, Charophyta, Ascomycota, Chytridiomycota, Chlorophyta and Bryophyta) were exclusive for deep-sea water samples (*Fig. 5b*).

**Figure 5 a, b:**
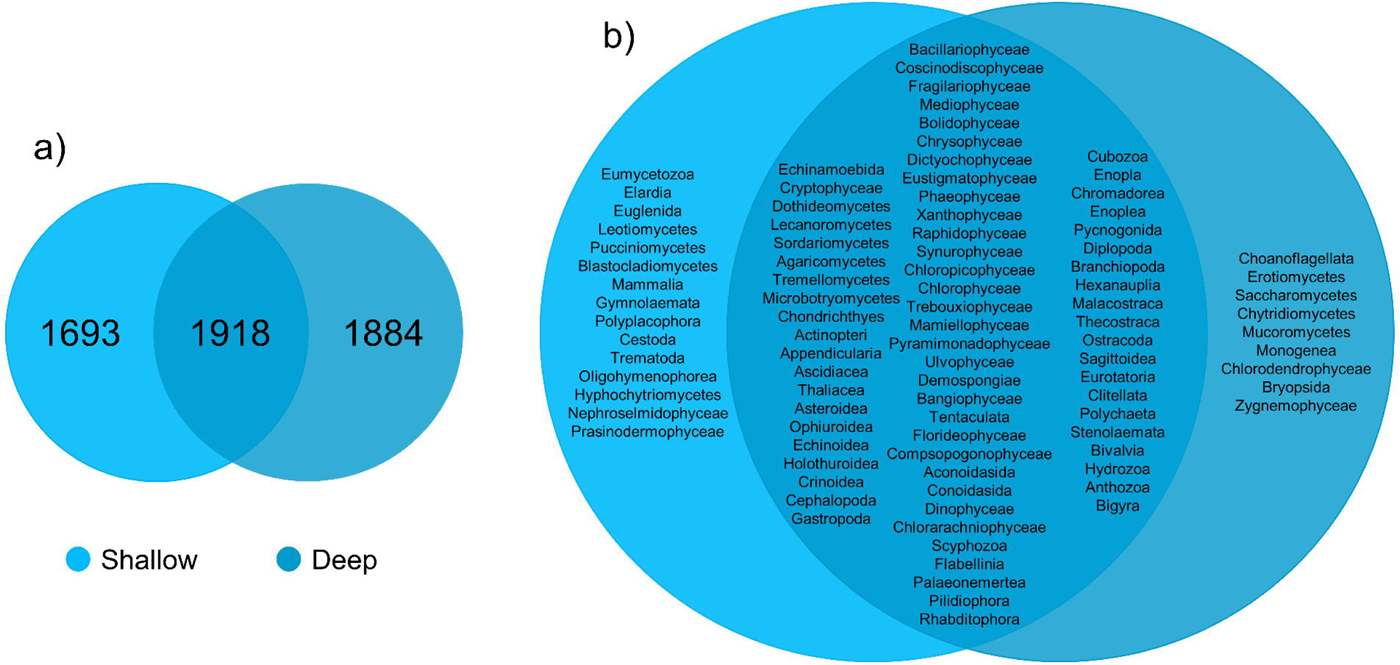
Beta Diversity. Venn diagrams represent the distribution of total OTUs (a) and shared and exclusive taxonomic classes per depth (b).

The nMDS ordination analyses categorized per biogeographic regions showed a concentration of the Northern Gulf of California sea water samples (*Fig. 6a*), but the ANOSIM test was not statistically significant (R = 0.11, *p* = 0.129, *Table S7*). A separation of shallow, mid-water depth, and deep-water samples was observed (*Fig. 6b*), where the shallow seawater samples were like each other (homogeneous). In contrast, samples from mid-water depth sites were more heterogeneous than the shallower sea water samples, but less contrasting than sea water samples from the deep-water layer. The ANOSIM test was statistically significant between depths (R =0.291, *p* = 0.001, *Table S7*). Regarding the water masses, the Tropical Surface Water (TSW) it is the most influential between shallow and mid-water depth sites, and where sampling sites grouped together independently of the region. The Gulf of California Water (GCW) basically influence the shallow and mid-water depth sites from the North region and one from the Central region (sc1). The Pacific intermediate Water (PIW) and the Subtropical Subsurface Water (StSsW) slightly separate the mid-water depth samples from the rest (*Fig. 6c*). However, ANOSIM analysis was not significant between water mass (R = 0.046, *p* = 0.3). The stress value (0.14) reflected a high ordination of the sampling sites. The ANOSIM test including biogeographic region, depth and water mass was statistically significant (R = 0.36, *p* = 0.002). The eukaryotic community structure based on the presence-absence of species at a global level was not statistically significant different, although did show a significant statistical difference per latitudinal region in the Gulf of California (PERMANOVA df = 29, Pseudo-F = 1.6467, *p* = 0.0002, *Table S8*). We showed evidence of significant differences in latitudinal regions after the pairwise comparison: North-Central (PERMANOVA df = 28, t = 1.3235, *p* = 0.002), North-South (PERMANOVA df = 8, t = 1.2196, *p* = 0.042), and Central-South (PERMANOVA df = 22, t = 1.2586, *p* = 0.0044) (*Fig. 6a*). We also found statistically significant differences comparing between shallow and deep-sea water depth samples (PERMANOVA df = 29, Pseudo-F = 1.3647, *p* = 0.007) (*Fig. 6b*). We only found statistically significant differences in depths: Shallow-Mid-water depth pairwise comparison (PERMANOVA df = 27, t = 1.3503, *p* = 0.00*2*). OTUs collected in different sea water mass exhibit a significant statistical difference (PERMANOVA df = 28, Pseudo-F = 1.2656, *p* = 0.006) (*Fig. 6b*). Following the pairwise comparison, we found significant differences in OTUS recorded from seawater samples collected in different sea water masses: GCW-TSW (PERMANOVA df = 27, t = 1.3067, *p* = 0.002), GCW-PIW: (PERMANOVA df = 8, t = 1.2492, *p* = 0.024) (*Fig. 6c*).

**Figure 6 a, b, c:**
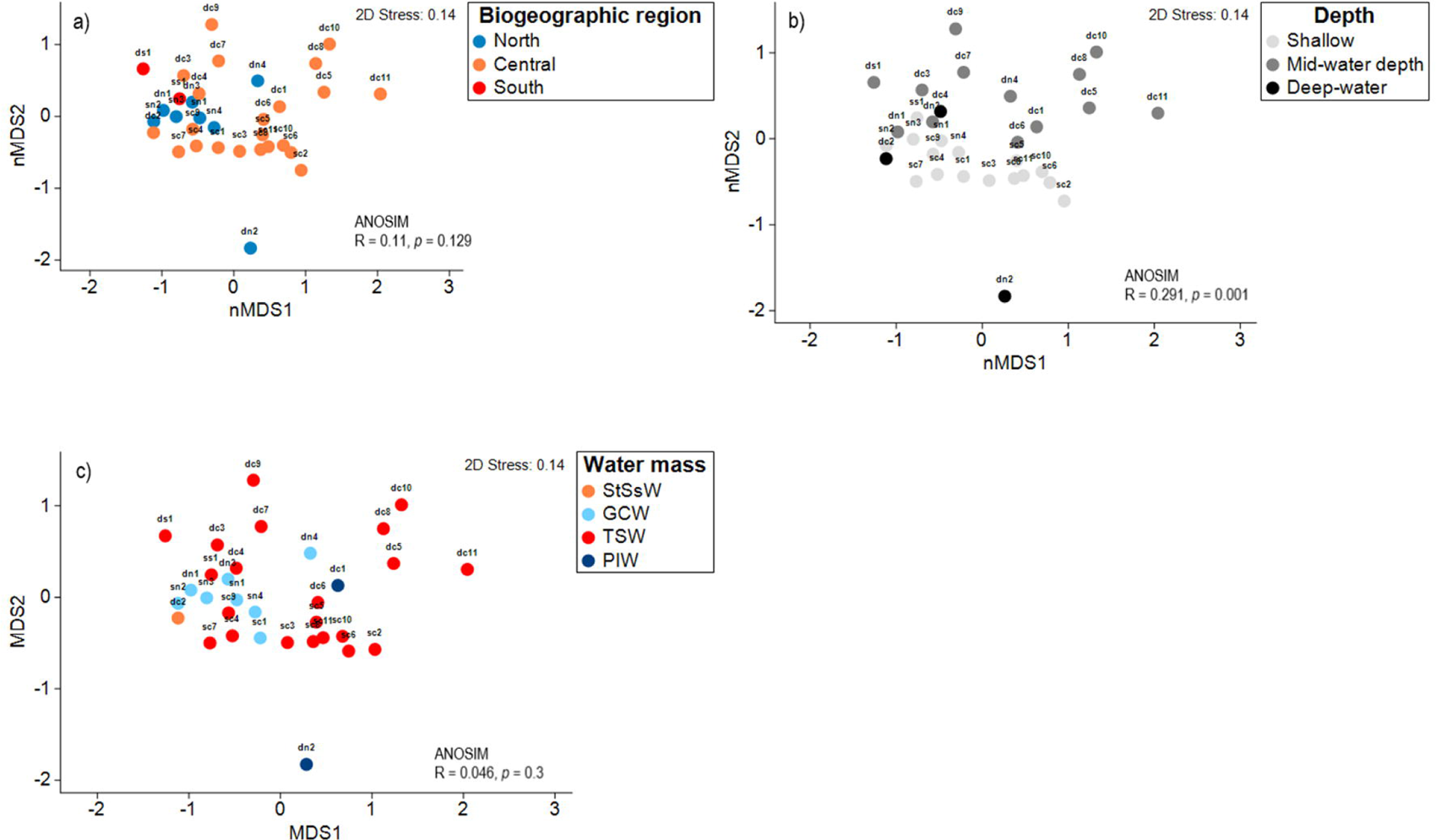
Eukaryotic Community Structure. Non-metric multidimensional scaling (nMDS) ordination analysis showing the OTUs community structure differences between three latitudinal regions in the Gulf of California (a), three seawater sampling depths (b), and four sea water masses from the Gulf of California (c).

The SIMPER analysis of species relative abundance richness showed that the differences found between different column water depths were related to taxa found in the shallow zone within the phyla Mollusca, Arthropoda, and several taxonomically unassigned OTUs. The difference between the North and Central latitudinal regions was mainly due to contribution of taxa associated with the deep zone within the phyla Chordata, Arthropoda, and Bacillariophyta where more than half of the OTUs observed classified as not assigned (*Table S9*). Between the Northern and Southern regions, the most important contribution was due to taxa associated with distinct sea water masses (Cnidaria, Rhodophyta, Echinodermata, and Arthropoda) and taxa from different sea water column depths (Arthropoda, Bacillariophyta, and three Not assigned OTUs).In the case of Central and South region comparison, the contribution to differentiation was related to taxa associated with both depth and sea water masses in the Central region (Cnidaria, Arthropoda, Rhodophyta, and two not taxonomically assigned OTUs). Finally, the Spearman rank correlation test (*Table S10*) suggested a poor relationship between column water depth and OTUs richness (R = -0.161), and the coefficient test result was not statistically significant different (R^2^ = 0.26, *p* = 0.377) (*Fig. 7*, *Table 10*).

**Figure 7:**
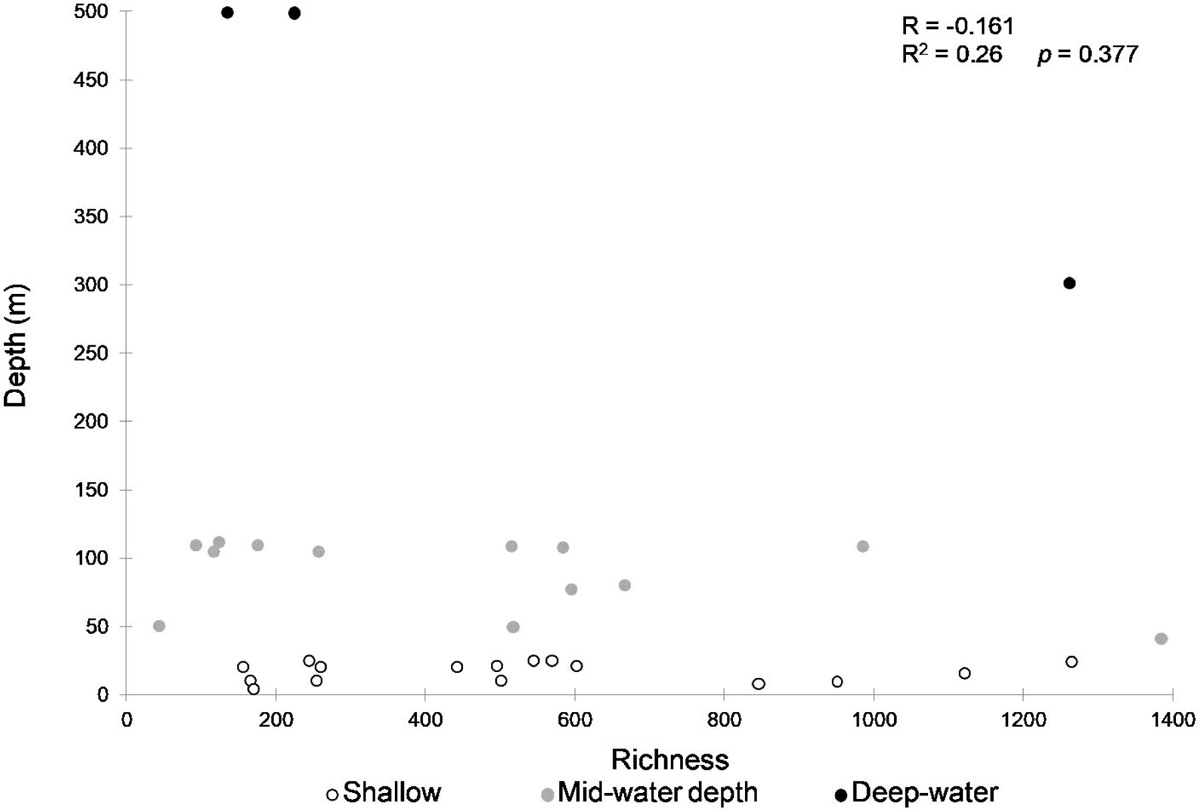
Relationship between Depth and OTUs richness. Scatter plot of Spearmańs rank correlation (R) and coefficient (R^2^) between depth and OTUs richness for the study area (95% CI).

## Discussion

We demonstrated that mid-water depths and deep-water samples showed similar levels of mean OTUs richness than shallow layers at the same sites and found evidence supporting the view that the communities found in the deeper layers were statistically distinct at the lowest taxonomic levels from their shallow layer counterparts. About a third of all the OTUs were exclusive from the deep-sea water samples and substantial turnover of OTUs was observed between the shallow and deeper sea water layers. Our results from the Gulf of California do not support the observed trend of decreasing marine biodiversity with depth (Costello & Chaudhary, 2017). These observations are in line with other studies along a similar vertically depth gradient comparing data from trawl surveys (Piacenza et al., 2015) and photo transects (Stefanoudis et al., 2019) reporting that biodiversity of the mid-water depth layer and the deep-sea is richer and more distinct than previously assumed. Other reports employing eDNA metabarcoding using universal primers also have detected significant vertical changes in the community composition of eukaryotes comparable to our study (Zhang et al., 2020; Govindarajan et al., 2021; Cote et al., 2023; Hoban et al., 2023). Our results imply that the deep refugia hypothesis could apply to only about a third of all the lower-level taxa identified that are shared between shallow and deep-sea water samples, while most OTUs could be considered exclusive from either shallow or deep-sea water layers.

The vertical transition in the biotic communities along a depth gradient could be explained by changes in temperature and availability of light and nutrients between the upper (30-60 m) and lower mid-water depth zones (60-150 m) (Lesser et al., 2019; Slattery et al., 2024). This vertical change is characterized by a sharp change in foundational species from photo autotrophic (hard corals and macroalgae) to heterotrophic taxa (sponges and azooxanthellate octocorals) feeding on large (>2 mm) zooplankton that seems more abundant in the mid-water depth layer (Andradi-Brown et al., 2017; Lesser et al., 2019). Fish species assemblages in the mid-water depth layer also show lower abundance of herbivores and higher population biomass (Stefanoudis et al., 2019; Rocha et al., 2018). Additionally, there is a vertical gradual transition to the deep sea represented by rariphotic ecosystems with unique species assemblages located between 150-300 m that are also different from the mid-water depth biota (Baldwin et al., 2018).

Marine biodiversity assessments have traditionally considered the ocean on a two-dimension scale with little focus on depth (Jacquemont et al., 2024). The advent of techniques like eDNA metabarcoding from seawater samples collected remotely opens new possibilities for characterizing marine biodiversity from deep ecosystems that were previously logistically inaccessible and thus lacked enough data. The finding that mid-water depths and deep-water zones are as OTUs diverse as shallow sea water OTUs assemblages and yet quite distinct in their species composition has some important implications for management and conservation. Based on a principle of complementarity, our study supports the view that marine protected areas (MPAs) and other effective area-based conservation measure should extend to include the biodiversity present in mid-water depth and deep-water layers to maximize protection of taxonomic and functional diversity of marine ecosystems against human impacts (Robison, 2009; Loiseau et al., 2022; Jacquemont et al., 2024). Even when some of the same species are present between the shallow and deep sea zones likely due to diel vertical migration (Ambriz-Arreola et al., 2017), the ecological and genetic connectivity between them could be limited, at least for some species that do not migrate of when migrate do not cross distinct vertical ecosystem (Palomares-García et al., 2013; Loya et al., 2016; Bongaerts et al., 2017). Copepods for example do not migrate daily maintaining maximum densities at 50 m depth independently of the time in the circadian cycle (Palomares-García et al., 2013). In contrast, krill migrates from surface to mid-water depths every day explaining sharing of species in different vertical habitats (Ambriz-Arreola et al., 2017). Some evidence suggest that ecosystems from the mid-water depth zone could act as a refugia for some key or foundational species (and the benthic communities associated with them) against stronger climatic impacts evident in shallower sites (Giraldo-Ospina et al., 2020) and being more resilient to local extinctions (del Monte-Luna et al., 2023). Extending the bathymetric range on which MPAs are placed and providing protection for the entire water column would also benefit a third of all the taxa that was not present in shallow layers and was exclusive from the deeper layers of the Gulf of California. Recent studies suggest that some of these taxa exclusive from deep waters are threatened with extinction, but many of these taxa are data deficient (Finucci et al., 2024).

Marine species closely track shifting isotherms due their metabolic needs in both pelagic and benthic habitats due to climate change shifting biogeographic regions latitudinally at ∼70 km/decade towards higher latitudes but also to greater depths (Lenoir et al., 2020; Pinsky et al., 2020). This reorganization of marine biodiversity highlights the need to monitor climate-driven community restructuring *in-situ* at different ocean depths. Projected range shifts based on climate velocities are faster in the deep ocean compared to the near surface, particularly for the mid-water depths biota (Brito-Morales et al., 2020). Since mid-water depth reefs are ecological relevant habitat for several economically profitable commercial species (Williams et al., 2019), the redistribution of species will have also economic impacts to the regional fishing sector (Gordo-Vilaseca et al., 2023; McClure et al., 2023) as has been documented in the Gulf of California (Gilly et al., 2022). With the availability of marine biodiversity data from the deep ocean, three-dimensional (3D) spatial prioritization analyses could be conducted and likely will become more common in the near future (Venegas-Li et al., 2018), and could incorporate climate velocities to identify climate refugia within present and future MPAs (Brito-Morales et al., 2022). Available data supports a strong ecological and biogeochemical connectivity between the pelagic (sea water column) and benthic mid-water depths and deep-water environments, but more observation evidence needs to be collected regarding the influence of vertically stratified management, particularly on oceanic MPAs (O’Leary and Roberts, 2018; Blanluet et al., 2023).

While depth layers showed the largest influence in shaping eukaryotic community composition, we found some evidence suggesting a role of sea water masses in the Gulf of California on species assemblage composition. Other studies have also shown distinct communities detected via eDNA metabarcoding associated with different sea water masses and driven by different planktonic organisms (Adams et al., 2023). The vertical distribution of multiple sea water masses layered in a long, narrow, and deep layers of the Gulf of California seem to contribute to a higher similarity of OTUs among sites located at similar depths and influenced by the same sea water mass. Each sea water mass represents a different habitat, characterized by multiple taxa responding to common environmental conditions (Lima-Mendez et al., 2015), in this case, the physical and chemical signatures that define it. The large difference observed between the community from the mid-water depth layer in the Northern Gulf of California (represented by sample dn2) from the rest of the samples could be attributed to the oceanographic isolation of the Northern Gulf of California by a series of islands, narrow channels, and sills from the central Gulf of California (Figure 1) that seems to separate the PIW water mass into two distinct regions as has been reported in zooplankton (Brinton et al., 1986; Quiroz-Martínez et al., 2023). The Northern Gulf of California is characterized by strong currents and complex topography that promotes year-round vertical mixing and primary productivity and high zooplankton biomass throughout the year (Salas-de-León et al., 2011). The use of eDNA metabarcoding for biodiversity monitoring beyond shallow coastal zones has several logistical challenges. While modeling and empirical studies have shown that the vertical distribution of eDNA often corresponds to the vertical location of the organismal source, sinking of eDNA has been proposed and implies that eDNA could be detected in the upper depth limit of any given taxa (Allan et al., 2021; Canals et al., 2021). This should be analyzed considering the role of natural diel vertical migrations on plankton and nekton that represent eDNA distribution in the water column (Easson et al., 2020; Canals et al., 2021). A decline of eDNA concentration in function of depth highlights the need of larger seawater volumes of filtered water and higher number of replicates are needed to improve the detection of pelagic biodiversity (McClenaghan et al., 2020; Govindarajan et al., 2022).

In our study, the proportion of taxonomically unassigned taxa and the slopes of the species accumulation curves indicate a larger sampling effort is required, particularly for the deeper layers, including broader geographic coverage, sampling, and sequencing efforts (both sequencing depth per sample and complementing DNA sequence databases), to comprehensively describe biodiversity.

## Acknowledgements

We thank Dr. Carlos Robinson for allowing us to collect samples during the 2018 CAPEGOLCA cruise. We thank the logistical support from the crew of the diving vessel Quino El Guardian.

This paper is part of the PhD research of PCE in Biosistemática, Ecología y Manejo de Recursos Naturales y Agrícolas (BEMARENA) postgraduate program at the University of Guadalajara.

She is also the recipient of a PhD scholarship granted by the CONAHCYT (CVU 268703). We thank Alma Paola Rodríguez-Troncoso and Camila Mac Loughlin for help filtering water samples.

