## SUpplemental Figure for "eDNA metabarcoding shows highly diverse but distinct shallow, mid-water, and deep-water eukaryotic communities within a marine biodiversity hotspot"

**Figure S1**

Tittle: Species Accumulation Curves

Legend: Chao1 cumulative richness curves per vertical layers: all eDNA sea water samples (a), shallow samples (b), and deep samples (mid-water and deep-water) (c) layers in the Gulf of California. None of the curves reached the asymptote suggesting sample size is still insufficient to capture the entire biodiversity present in the samples.
