## Supplementary figures and images for "eDNA metabarcoding shows highly diverse but distinct shallow, mid-water, and deep-water eukaryotic communities within a marine biodiversity hotspot"

### Supplemental Figure S1

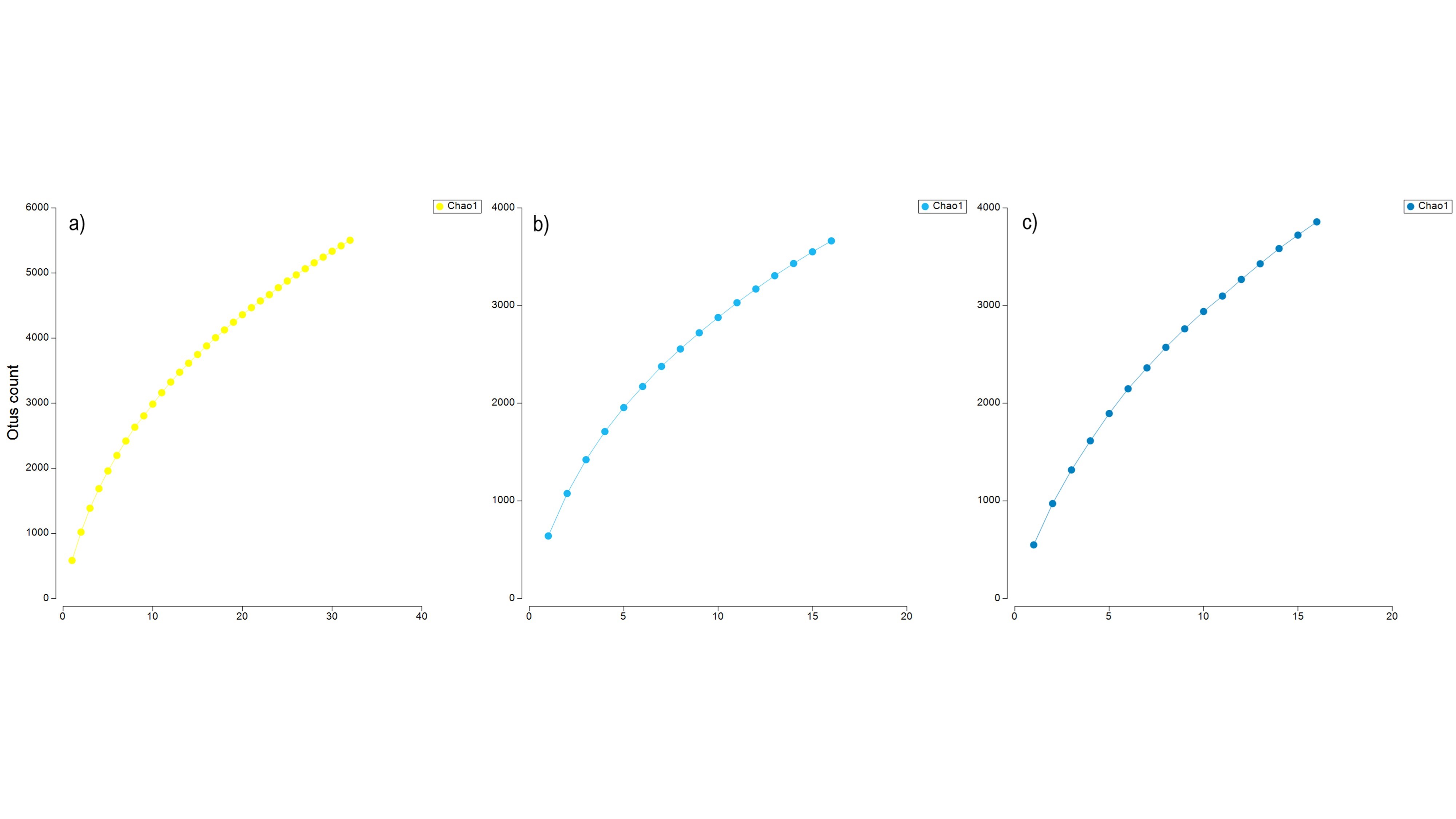
